# Novel triplex nucleic acid lateral flow immuno-assay (NALFIA) for rapid detection of Nipah virus, Middle East respiratory syndrome coronavirus and Reston ebolavirus

**DOI:** 10.1101/2022.10.06.511237

**Authors:** Santhalembi Chingtham, Diwakar D Kulkarni, TS Sumi, Anamika Mishra, Atul K Pateriya, Vijendra Pal Singh, Ashwin Ashok Raut

## Abstract

We report the development of the first triplex Nucleic Acid Lateral Flow Assay (NALFIA) for detection of genomes of Nipah virus (NiV), Middle East respiratory syndrome coronavirus (MERS-CoV) and Reston ebolavirus (REBOV), intended for screening of bats as well as other hosts and reservoirs of these three viruses. Our triplex NALFIA is a two-step assay format wherein the target nucleic acid in sample is first amplified using tagged primers, and the tagged ds DNA amplicons are captured by immobilized antibodies on NALFIA device resulting to signal development from binding of streptavidin-colloidal gold conjugate to biotin tag on the captured amplicons. Triplex amplification of *N* gene of NiV, *UpE* gene of MERS-CoV, and *Vp40* gene of REBOV was optimized using primers adapted from validated real-time RT-PCR assays of previous studies and the compatible combinations of hapten-labels and antibodies for triplex NALFIA device were identified. Digoxigenin, rhodamine red and alexa fluor 488 were identified as suitable 5’ labels on forward primers. The lowest copy number detected by the triplex NALFIA with 2 μl of triplex RT-PCR product were up to 8.21e4 for NiV *N* target, 7.09e1 for MERS-CoV *UpE* target, and 1.83e4 for REBOV *Vp40* target. Using simulated samples and Taqman real-time RT-PCR as standard, the sensitivity and positive predictive values were found to be 100% for MERS-CoV *UpE* and REBOV *Vp40* targets and 91% for NiV *N* target while the specificity and negative predictive values were 100% for MERS-CoV *UpE* targets and REBOV *Vp40*, and 93.3% for NiV *N* target.

## Introduction

The devastating implications of emerging zoonotic infectious diseases are well recognized. Nipah virus (NiV), Middle-East respiratory syndrome coronavirus (MERS-CoV) and ebolaviruses are emerging zoonotic viruses, which account for high case fatality rates in human, incur economic losses and affect international travel and animal trade due to livestock involvement (1, 2, 3). Reston ebolavirus (REBOV) is non-lethal to humans (4, 5), however, high pathogenicity of REBOV in non-human primates, the non-clinical susceptibility of domestic pig and human to REBOV infection, and the prevalence of REBOV in Asia unlike other ebolaviruses that are confined in a particular geographical area outside Asia (6, 7, 8, 9) suggests the possibility of emergence of mutant REBOV with increased lethality in human and livestock. Bats are reservoirs of henipaviruses, coronaviruses and ebolaviruses, although direct or indirect involvement of bats in MERS-CoV transmission is yet to be identified (10, 11, 12, 13, 14, 15). Therefore, it is imperative to screen reservoir bats for assessment of potential health risks and threats imposed by NiV, MERS-CoV and REBOV in Asia. The present study highlights the development of a triplex one-step RT-PCR-based triplex nucleic acid lateral flow immuno-assay (NALFIA) for rapid and simultaneous screening of samples, particularly of bats, for NiV, MERS-CoV and REBOV by targeting the viral RNA. Multiplexing allows the use of a single sample for screening of multiple pathogens in a single test; hence making the assay rapid, cost-effective and also reducing the need for multiple sample processing from an individual animal to test multiple pathogens, a consideration vital to screening wildlife (bat) samples.

For detection of viral infections, molecular tests are superior to non-nucleic acid diagnostic tests such as serology and microscopy in terms of sensitivity as well as specificity (16). Conventional PCR-based tests are sensitive; however, end-point detection of PCR ampliconsby agarose gel electrophoresis is complex considering the operation, hazard, time and cost. While the real-time PCR system does not need an additional post-run wet lab procedure, its downsides are high costs and sophistication that limit its usage at low resource settings. An efficient alternative read-out system is the NALFIA, which is a sub-type of the lateral flow assay (LFA) (17). NALFIA uses immobilized antibody to capture double stranded DNA amplicons through a tag on one of the oligo primers. Another tag is incorporated into the amplicon, through one of the primers, which has affinity for the detector conjugate, the interaction of which creates the signal on the test line and the conjugate control line. Therefore, NALFIA can also be interpreted as a two-step assay format wherein the target nucleic acid in sample is first amplified, followed by examination of amplicons on the LFA device. NALFIA device offers an augmented ease in operation of the PCR setup and a very rapid and safe detection of the amplified products of viral genome. Another advantage of NALFIA device is that it provides an improved specificity and sensitivity over AGE in detection of amplified products owing to its dual label and antibody-based interactions. PCR-based monoplex or multiplex NALFIA for the detection of DNA targets have been reported in previous studies (16, 18, 19, 20). And recently, isothermal assay-based multiplex NALFIA has also been developed for detection of SARS-CoV2 (21, 22, 23).

We report the development of a triplex one-step RT-PCR-based NALFIA with three test lines and a conjugate control line for simultaneous detection of NiV, MERS-CoV and REBOV RNA targets. Oligonucleotide primers used in the present work were adapted from validated real-time RT-PCR assays developed in previous studies and extensively used for diagnosis of NiV, MERS-CoV and REBOV infections (24, 25, 26). However, we optimized the triplex RT-PCR amplification conditions and the triplex hapten label combinations for the NALFIA device for the three selected targets. The gene targets for NiV, MERS-CoV and REBOV detection employed in our assay were Nucleoprotein (*N*), upstream E (*UpE*) and matrix (*Vp40*), respectively. The triplex NALFIA was standardized using IVT RNAs generated from an in-house designed synthetic DNA template. A schematic representation of the components of our triplex NALFIA is shown in (Fig 1). From this point onwards in our current text, our triplex one-step RT-PCR based triplex NALFIA will be referred to as triplex NALFIA.

**Fig 1:**
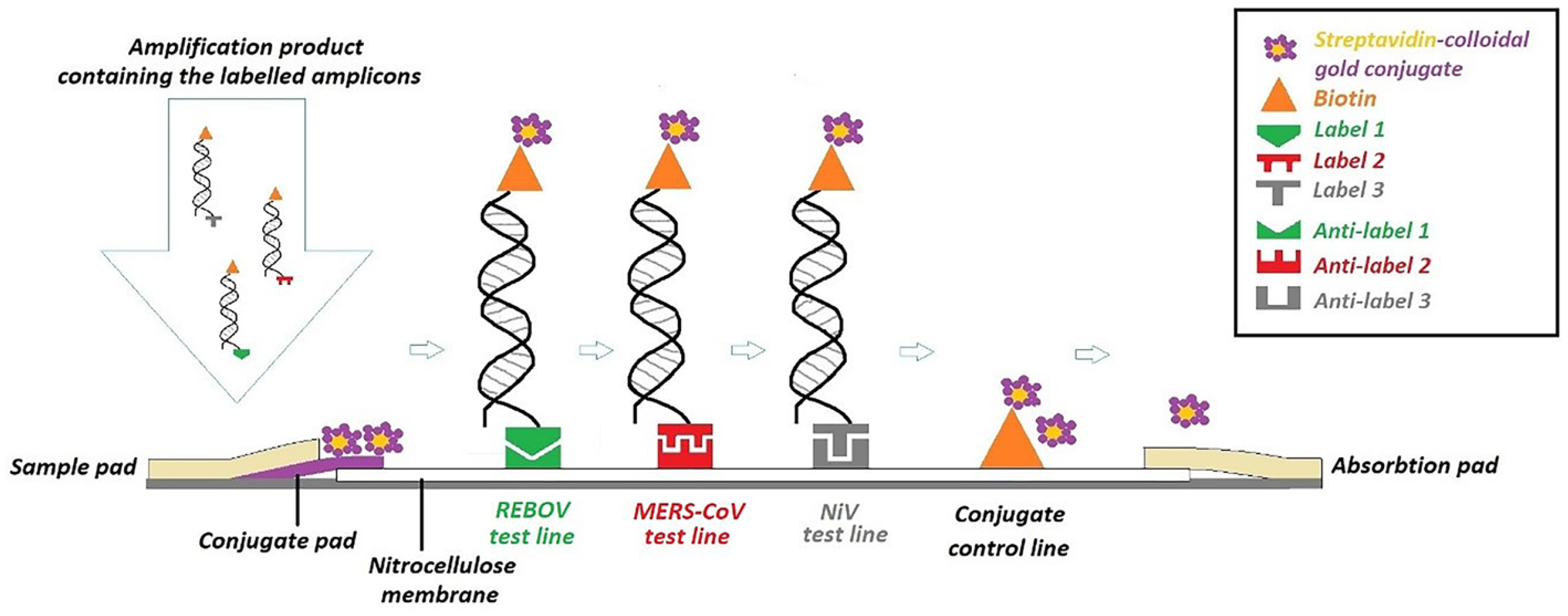
Schematic diagram of triplex NALFIA. 5’ labelled duplex target amplicons are generated by RT-PCR using forward and reverse primers with 5’ tags. The labelled amplicons are then captured through the tags on the forward primer by corresponding anti-tag/label antibody, which is the test line on the LFA device, while the biotin labelled on the reverse primer helps capture the detector system i.e., stretavidin-colloidal gold conjugate and produces the signal on the test line in presence of specific amplification. Label 1 - Alexa fluor 488, Label 2 - Rhodamine Red, Label 3 - Digoxigenin.

## Materials and Methods

### Primer sequences for NiV, MERS-CoV and REBOV

Nucleoprotein (*N*) gene of NiV, upstream E *(UpE)* gene of MERS-CoV and matrix *(Vp40)* gene of REBOV were selected as targets for detection as these genes are sensitive diagnostic markers. Based on the diagnostic recommendation by referral bodies (WHO and WOAH) the forward and reverse primers targeting these genes were selected from published literatures as listed in (Table 1).

**Table 1:**
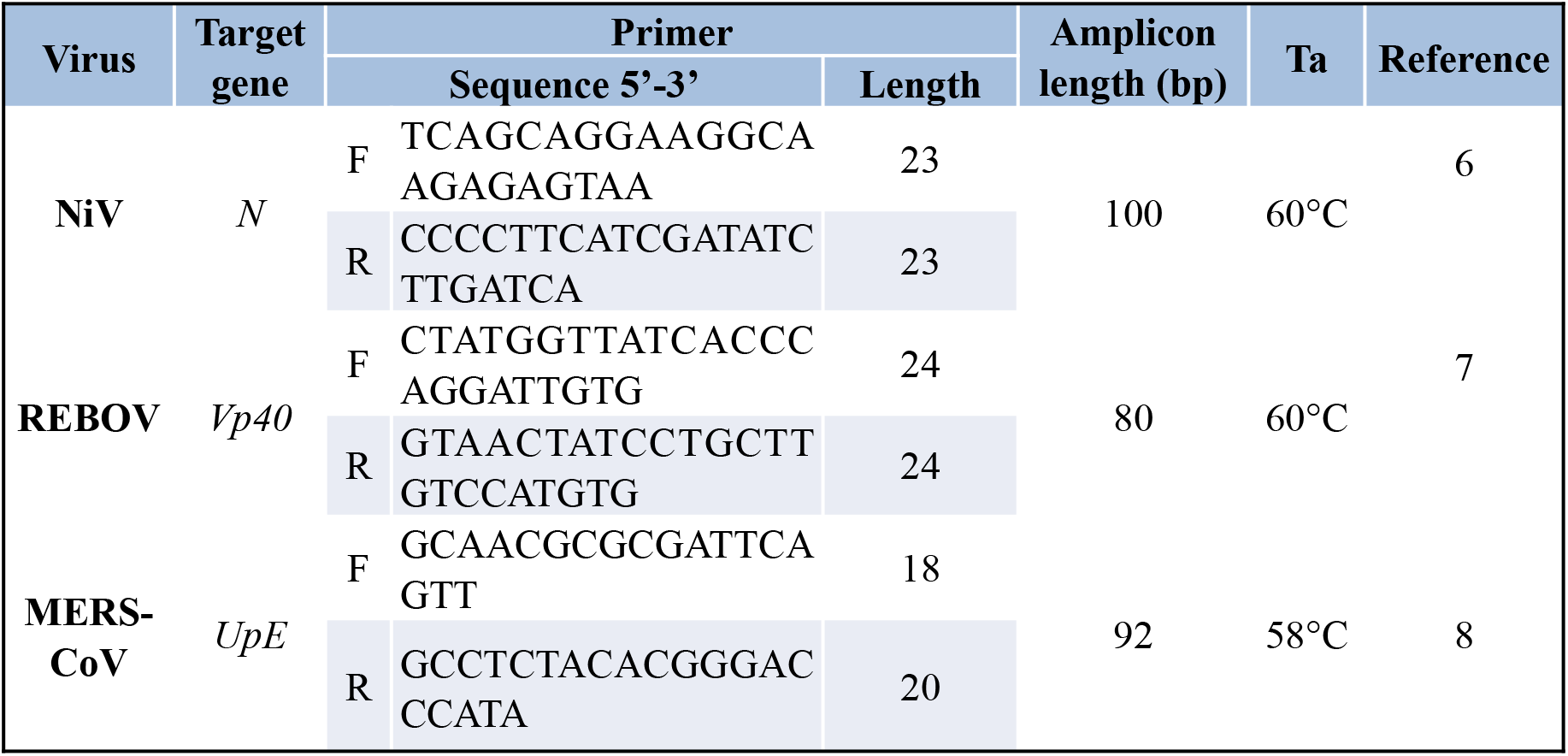
Details of primers used in the development of triplex NALFIA

### Template design and IVT RNA preparation

The nucleotide sequences of the PCR targets of NiV *N* (100 bp), MERS-CoV *UpE* (92 bp) and REBOV *Vp40* (80 bp) were downloaded from NCBI genbank and were constructed *in silico* in tandem using Editseq of DNAstar Lasergene software. The construct was synthesized by Genscript, USA in pBluescript II SK (+) plasmid (Fig 2).

**Fig 2:**
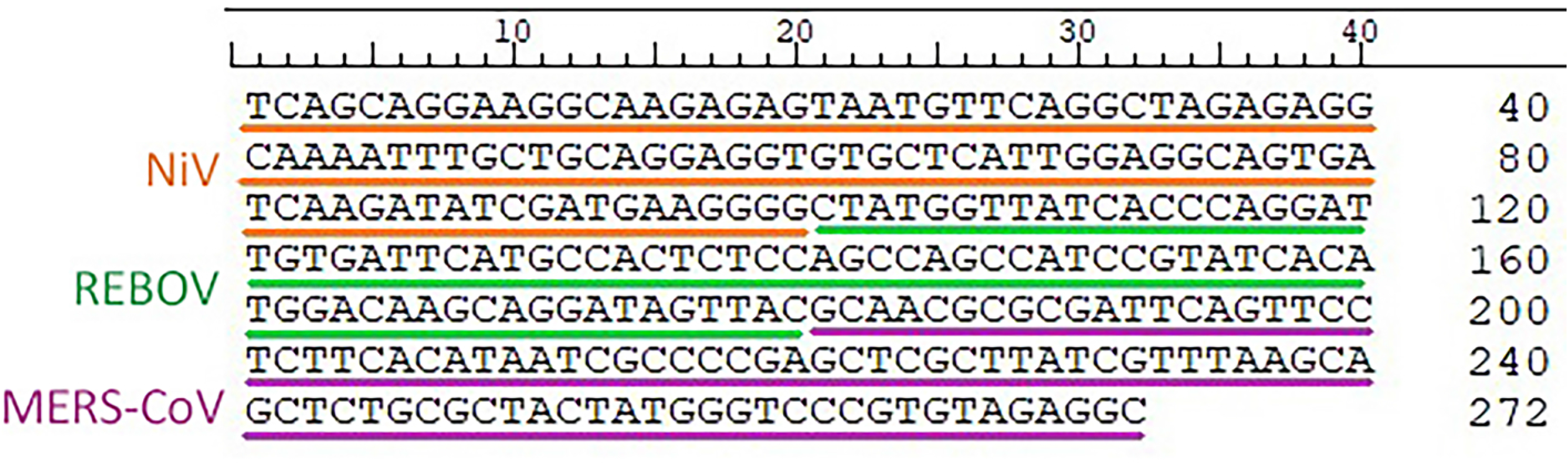
Schematic diagram of NALFIA triplex layout with coated lines, signal development and result interpretation. Signals at test line 1 and control line indicates a positive test for REBOV and negative for NiV and MERS-CoV. Signals at test line 2 and control line indicates a positive test for MERS-CoV and negative for REBOV and NiV. Signals at test line 3 and control line indicates a positive test for NiV and negative for MERS-CoV and REBOV. No signal at conjugate control line shows the test is invalid.

Since the fusion target DNA construct would possibly generate multiple non-specific amplicons when used as template with triplex primer cocktail owing to multiple combinations of forward and reverse primer binding sites present in tandem, it was necessary to separate the targets from the fused construct to make it functional as discrete templates for triplex primer cocktail. Therefore, individual target sequences were PCR amplified and the amplicons were sub-cloned into separate pGEM-T Easy vector (Promega) at TA cloning site. The resulting clones were then confirmed by restriction enzyme digestion – NiV *N*-pGEM-T Easy plasmid by *Pst I*, REBOV *Vp40-* pGEM-T Easy plasmid by *EcoR I and* MERS-CoV *UpE*-pGEM-T Easy plasmid by *Ava I*.

For *in vitro* transcription, the subcloned plasmids NiV *N*-pGEM-T Easy, REBOV *Vp40-* pGEM-T Easy and MERS-CoV *UpE*-pGEM-T Easy were first linearized by *Sal I* enzyme (Promega) creating a 5’overhang. RNA of NiV *N*, REBOV *Vp40* and MERS-CoV *UpE* targets were transcribed from 600 ng of linearized and purified plasmids using the TranscriptAid T7 High Yield Transcription kit (Thermo Scientific). The IVT mastermix was set up according to the manufacturer instructions and the reaction was performed at 37°C for 6-7 hr. The estimated transcript lengths were 166 nt (NiV) 146 nt (REBOV) and 158 nt (MERS-CoV). The IVT products were treated with 3U of Turbo DNase I enzyme (Ambion) and were purified 2-3 times by Trizol LS (Ambion) to completely remove the carry over plasmid DNA template. The IVT RNAs were quantitated by Qubit fluorometer using Quant-it RNA BR reagent (Molecular Probes) and stored at −80°C in small aliquots until further use. RNA copy number was calculated using online software, NEBioCalculator by New England Biolabs (https://nebiocalculator.neb.com) and the formula: RNA copy number = moles of RNA x 6.022 x10e23 (27).

### Optimization of triplex amplification condition

A triplex primer stock cocktail was prepared by mixing the NiV *N*, MERS-CoV *UpE* and REBOV *Vp40* specific forward and reverse primers in equimolar ratio and was diluted in a series to yield 8 different equimolar primer cocktails of 1.2 μM, 0.8 μM, 0.6 μM, 0.4 μM, 0.3 μM, 0.2 μM, 0.15 μM and 0.1 μM composite concentration. Each cocktail concentration was subjected to PCR amplification of individual sub-cloned DNA target plasmid templates, 10.5 pg of NiV *N*-pGEM-T Easy, 127 pg of REBOV *Vp40-pGEM-T* Easy and 7.7 pg of MERS-CoV *UpE-p* GEM-T Easy using 2X Verso hot-start PCR buffer (Thermo Scientific). The reaction was set up in 15 μl final volume and the thermal cycling was as follows: hot-start and initial denaturation at 95°C for 15 min, 35 cycles of denaturation at 95°C for 15 sec, annealing at 58°C (MERS-CoV target) and 60°C (NiV and REBOV targets) for 30 sec and extension at 72°C for 15 sec followed by final extension at 72°C for 5 min and 4°C hold. Using the final optimum composite primer concentration of 0.6 μM, the annealing temperature of the triplex amplification was optimized by testing four different temperature - 48°C, 52°C, 56°C and 60°C in thermal cycler with veriflex block (ABI Veriti 96-Well Fast Thermal Cycler, Thermo Fisher). The reaction was set up in 15 μl final volume and the thermal cycling was as follows: hot-start and initial denaturation at 95°C for 15 min, 35 cycles of denaturation at 95°C for 15 sec, annealing at 48°C, 52°C, 54°C and 60°C for 30 sec and extension at 72°C for 15 sec, followed by final extension at 72°C for 5 min and 4°C hold. The triplex amplification was also observed for any non-specific interactions among primers (homodimers and heterodimers) and between primers and non-self templates.

### Confirmation of IVT RNA

The IVT RNA of NiV, MERS-CoV and REBOV were diluted to 2×10^-3^ and confirmed by their respective monoplex one-step RT-PCR with specific primers using the Verso 1-step RT-PCR kit (Thermo Scientific). The reaction was set up in 10 μl final volume with 100 nM each of unlabelled forward and reverse primers, separately for each target. The thermal cycling was as follows: RT step at 50°C for 15 min, hot-start and initial denaturation at 95°C for 15 min, 35 cycles of denaturation at 95°C for 15 sec, annealing at 60°C for 30 sec and extension at 72°C for 15 sec, followed by final extension at 72°C for 5 min and 4°C hold. IVT RNA products (2×10^-3^ dilution) were tested for absence of carry-over plasmid template by PCR using Taq 2X master mix (NEB). A negative PCR result would confirm the absence of carryover plasmid in the IVT product.

### Optimization of compatible combinations of hapten labels and antibodies for triplex NALFIA

For selective and specific detection of our targets, the forward primers were tagged at 5’ end with different hapten labels to be captured by their corresponding antibodies immobilized on the lateral flow nitrocellulose membrane. For generating colour signal, all reverse primers were tagged at 5’ end with biotin in order to bind with streptavidin-colloidal gold conjugate. Different hapten labels were initially tested with corresponding capture antibodies (Table 2) for sensitive and specific interactions between labels and capture antibodies in a preliminary batch of monopolex, duplex and triplex lateral flow nitrocellulose strips (data not shown). For further testing of label-antibody compatibility in multiplexing, a hexaplex lateral flow nitrocellulose strips were also fabricated and tested with labelled amplicons of PCR. Out of six label-antibody combinations, the best three combinations were selected and our final triplexed NALFIA device was fabricated. The order of antibody test lines and control line in hexaplex and triplex formats of lateral flow strips are displayed in (Table 3).

**Table 2:** Hapten labels for 5’ end-labelling of forward and reverse primers and their corresponding antibodies used in the optimization of multiplexing compatibility of hapten-antibody combination for the triplexed NALFIA device.

| Combination | 5' hapten label on forward primer | Antibody against 5' label on forward primer | 5' hapten label on reverse primer |
| --- | --- | --- | --- |
| 1 | Dinitrophenyl (DNP), IDT | Polyclonal anti-DNP ab, Molecular Probe | Biotin, Eurofins |
| 2 | Digoxigenin (DIG), Eurofins | Polyclonal anti-DIG ab, Pierce | Biotin, Eurofins |
| 3 | Texas Red, Eurofins | Polyclonal anti-texas red ab | Biotin, Eurofins |
| 4 | Fluorescein, Eurofins | Polyclonal anti-fluorescein ab, Molecular Probe and Monoclonal anti-fluorescein ab, Molecular Probe | Biotin, Eurofins |
| 5 | Rhodamine Red, Eurofins, IDT | Polyclonal anti-tetramethyl rhodamine ab, Molecular Probe | Biotin, Eurofins |
| 6 | Alexa fluor 488, Eurofins | Polyclonal anti-alexafluor 488 ab, Molecular Probe | Biotin, Eurofins |

**Table 3:** Layout of hexaplex and triplex NALFIA strips with specific order of the label antibodies as test lines

| Hexaplex |  | Triplex |  |
| --- | --- | --- | --- |
|  | *Absorbtion Pad* | *Absorbtion Pad* |  |
| **Control Line** | Conjugate control | Conjugate control | **Control Line** |
| **Test Line 1** | anti-DNP ab | anti-DIG ab | **Test Line 1** |
| **Test Line 2** | anti-DIG ab | anti- Rhodamine ab | **Test Line 2** |
| **Test Line 3** | anti-Fluorescein ab | anti-Alexa Fluor 488 ab | **Test Line 3** |
| **Test Line 4** | anti-Texas Red ab | *Sample pad/Conjugate Pad* |  |
| **Test Line 5** | anti-Alexa fluor 488 ab |  |  |
| **Test Line 6** | anti-Rhodamine ab |  |  |
|  | *Sample pad/Conjugate Pad* |  |  |

DNP - Dinitrophenyl; DIG - Digoxigenin

### Nucleic acid lateral flow immuno-assay (NALFIA) device

NALFIA device with three test lines and a control line were custom manufactured from Ubio Biotechnology Pvt. Ltd., Cochin, (India) in a batch method. Stripes of 0.5 mm width of each capture antibody were applied at a rate of 0.11 μl/mm as 3 consecutive test lines at a concentration of 1 mg/ml in coating buffer on nitrocellulose membrane of 0.47 mm thickness and 0.22 μm pore size. Biotin-BSA was applied as the conjugate control line of 0.5 mm width at 0.75 mg/ml. Conjugate pad made of 5 mm wide glass fiber was impregnated with streptavidin-colloidal gold conjugate (10 μg/ml) by soaking in 1:2 dilution of conjugate solution, prepared at 1:1 ratio in conjugation buffer. Sample pad was 18 mm wide glass fiber while absorption pad was 20 mm cellulose fiber. 3.2 mm wide individual strips were cut after manual lamination of the components followed by manual housing (Polypropylene 6.9 cm x 2 cm, TV Plastics). All handling was performed at ≤10% humidity and 30°C room temperature.The schematic representation of the NALFIA device with interpretation is shown in (Fig 3).

**Fig 3:**
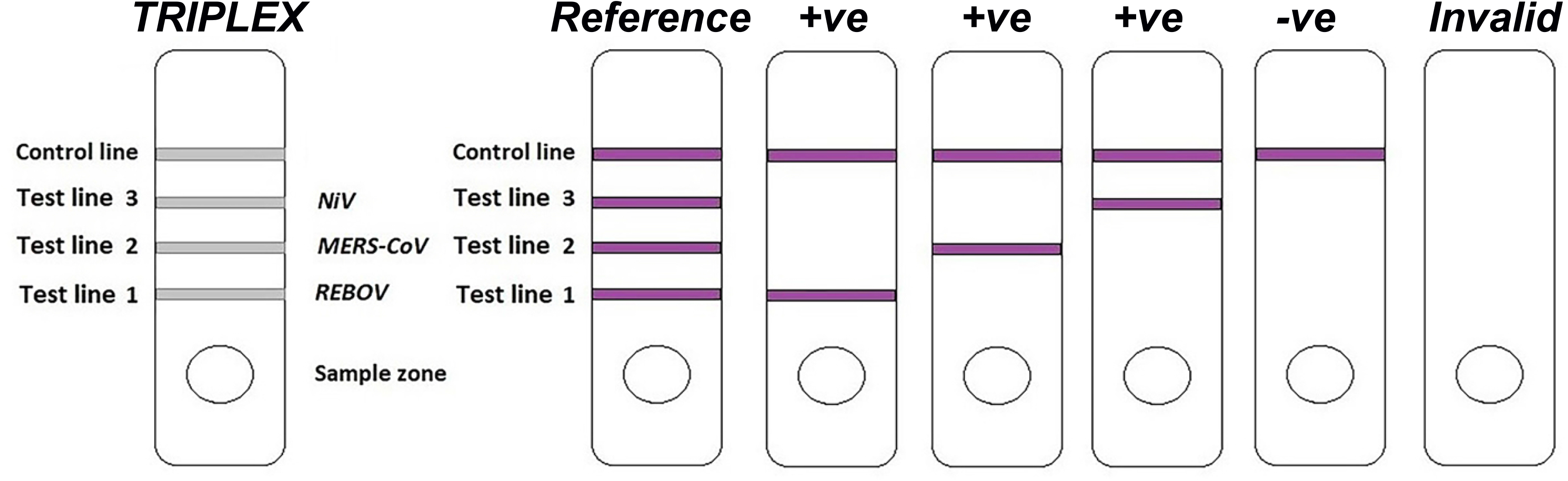
A 272 bp gene construct comprised of targets of NiV *N* (100 bp), REBOV *Vp40* (80 bp) and MERS-CoV *UpE* (92 bp) synthesized in tandem at EcoR I site in pGEM-T Easy vector (vector not included in the figure). NCBI Genbank accession numbers of the source of the target genes are EU620498.1 (NiV), AB050936.1 (REBOV), and JX869059.2 (MERS-CoV).

### Sensitivity of NALFIA device for labelled PCR amplicons

Two micro litres of undiluted labelled monoplex PCR products of each target and their log dilutions were added into sample pads of NALFIA devices, separately. The migration buffer (10 mM PBS, 1% BSA, 0.05% Tween 20, pH 7.5) was added drop wise after adding the PCR products until the conjugate front disappears. The band intensity in each device was subsequently observed within a signal development period up to 15 min. The bands on devices were compared with corresponding band gradation in 3% AGE of each target of 5 μl load volume.

### Determination of limit of detection (LOD) of Triplex NALFIA

10-point serial log dilutions (10^-1^ to 10^-10^) of quantitated stock IVT RNA template of NiV, MERS-CoV, and REBOV targets were prepared and each dilution of the three targets were subjected to amplification by triplex one-step RT-PCR. Using Verso 1-Step RT-PCR kit (Thermo Scientific), reactions of 13 μl final volume was set up using 0.6 μM of triplex primer cocktail in equimolar ratio of labelled forward and reverse primers of each target. The linearity of the amplification was visualised using 5 μl products in 3% agarose gel in 1x TAE buffer and the sensitivity or the limit of detection of the triplex NAFIA was evaluated using 2 μl of the product on the NALFIA device.

### Specificity of triplex NALFIA

The specificity of triplex NALFIA was evaluated by observing any non-specific signals arising out of interactions among the primers in the triplex cocktail, and between the triplex primers and the templates. The specificity of the triplex NALFIA was also analyzed by testing clinical samples negative for NiV, MERS-CoV and REBOV but positive for *Flaviviridae* (Japanese Encephalitis Virus, JEV; Classical swine fever virus, CSFV), *Paramyxoviridae* (New castle disease virus, NDV), *Coronaviridae* (Infectious bronchitis virus, IBV), *Orthomyxoviridae* (Avian influenza virus, AIV), *Picornaviridae* (Foot and mouth disease virus, FMDV), *Herpesviridae* (Duck plague virus, DPV), *Arteriviridae* (Porcine reproductive and respiratory syndrome virus, PRRSV), *Poxviridae* (Suipox virus, SPV).

### Evaluation of triplex NALFIA with samples spiked with IVT RNA and unspiked samples

For simulation of positive samples, an assorted group of samples comprising of sera, oropharyngeal swab, tissue and whole blood from bat *(Pteropus giganteus)* camel and pig that represents the target species and sample-source population of the three select viruses were spiked (S1) with 28 pg to 56 pg NiV *N* IVT RNA, 230 fg to 460 fg MERS-CoV *UpE* IVT RNA, and 5.5 pg to 11 pg REBOV *Vp40* IVT RNA at lysis step of RNA extraction. TaqMan RT-qPCR was performed to authenticate the samples. The unspiked and spiked samples were then subjected to amplification using the Verso 1-step RT-PCR kit (Thermo Scientific); the reaction was set up in 10 μl final volumes with 600 nM of labelled triplex primer cocktail and 1-3 μl of RNA template. The thermal cycling was as follows: RT step at 50°C for 15 min, hot-start and initial denaturation at 95°C for 15 min, 35 cycles of denaturation at 95°C for 15 sec, annealing at 60°C for 30 sec and extension at 72°C for 15 sec, followed by final extension at 72°C for 5 min and 4°C hold. For readout on NALFIA device, 2 μl of triplex RT-PCR product was loaded on the NALFIA devices and the results were read between 1-15 min. Using real-time RT-PCR as standard test, the sensitivity, specificity and positive and negative predictive values (PPV and NPV) for the triplex NALFIA were calculated (28).

### Repeatability of triplex NALFIA

The triplex-NALFIA was performed three times with two spiked samples for each virus target to demonstrate the assay repeatability.

## Results

### Confirmation of the sub-cloned NiV N-pGEM-T Easy, REBOV Vp40-pGEM-T Easy and MERS-CoV UpE-pGEM-T Easy plasmid

Digestion of the NiV *N*-pGEM-T Easy plasmid of size 3115 bp by *Pst I* enzyme released two fragments of sizes 3037 bp and 78 bp, as *Pst I* cuts at positions 54/50 of the insert and 88/92 of the vector. Digestion of the REBOV *Vp40-pGEM-T* Easy plasmid by *EcoR I* enzyme yielded a single linearized plasmid band of 3095 bp size, as EcoR I cuts the plasmid at a single site. Digestion of the sub-cloned MERS-CoV *UpE*-pGEM-T Easy plasmid by *Ava I* enzyme resulted a single linearized plasmid band of size 3107 bp, as *Ava I* cuts only the insert at 36/40. The digestion pattern of the plasmids revealed that the target inserts were appropriately subcloned in pGEM-T Easy vector.

### Optimization of primer cocktail concentration and the annealing temperature (Ta) for triplex amplification

The optimum equimolar composite concentration of triplex primer cocktail of NiV *N*, MERS-CoV *UpE* and REBOV *Vp40* targets was found to be 0.6 μM (Fig 4). There was significant reduction in amplification at concentrations below 0.6 μM, while concentration above 0.6 μM resulted in dimers or excess primers. Amplification of NiV *N* and REBOV *Vp40* was evident till 150 nM, while MERS-CoV *UpE* amplification was evident till 100 nM. The amplification was uniform for all three targets at all the four tested annealing temperatures (Fig 5). The optimized annealing temperature of the triplex PCR was 60°C because primer-dimer was observed to be least or none at 60°C.

**Fig 4:**
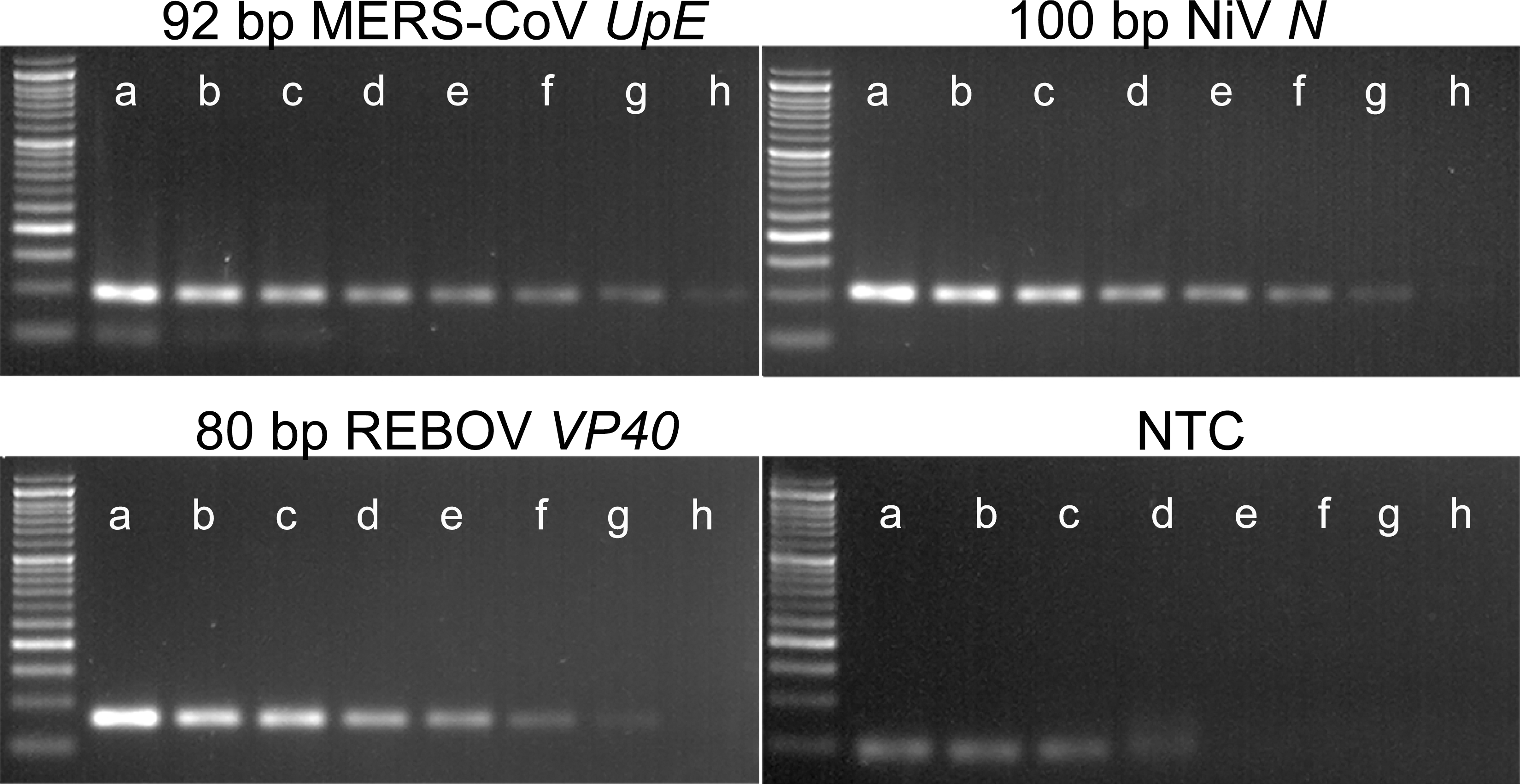
Optimization of triplex primer concentration for amplification. PCR amplification was analysed using different concentration of triplex primer cocktail comprised of forward and reverse primer sets of MERS-CoV, NiV and REBOV targets in equimolar ratio. Lane a - 1.2 μM, Lane b - 0.8 μM, Lane c - 0.6 μM, Lane d - 0.4 μM, Lane e - 0.3 μM, Lane f - 200 nM, Lane g - 150 nM, Lane h - 100 nM. Ladder - 50 bp.

**Fig 5:**
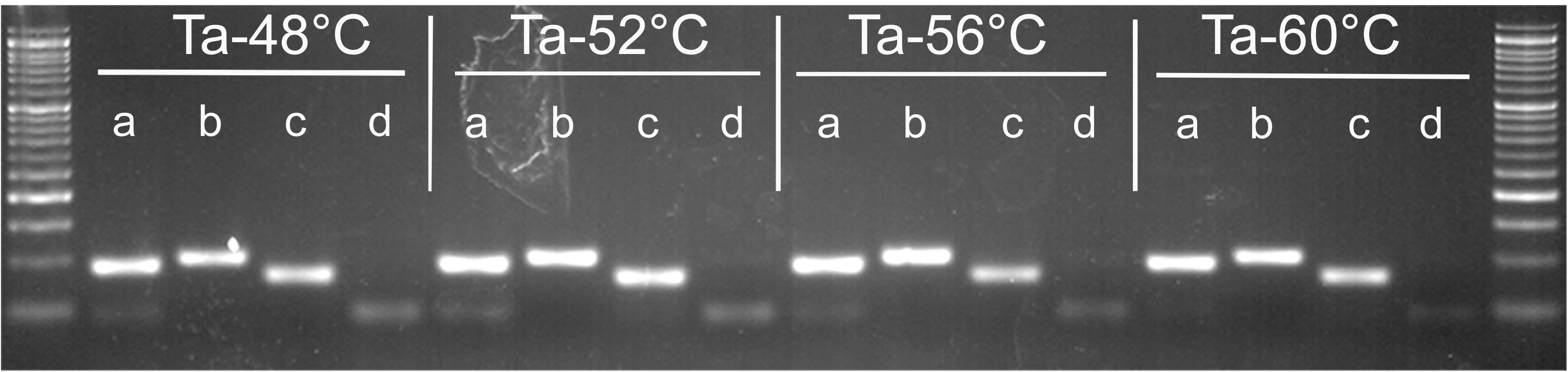
Optimization of annealing temperature for amplification. PCR optimisation using different annealing temperatures for triplex amplification using the optimised triplex primer cocktail. Lane a - MERS-CoV (92 bp), lane b - NiV (100 bp), lanec - REBOV (80 bp), and lane d - NTC. Ladder - 50 bp.

### In vitro transcription and confirmation

The *in vitro* transcription yielded NiV *N*, MERS-CoV *UpE* and REBOV *Vp40* RNA transcripts of approximately 166 nt, 158 ntand146 nt, respectively. The PCR of NiV *N*, MERS-CoV *UpE* and REBOV *Vp40* resulted in no amplification indicating the absence of carryover DNA in the IVT product (Fig 6). The one-step RT-PCR of IVT RNA gave specific amplicons indicating the amplification exclusively from IVT RNA template (Fig 6).

**Fig 6:**
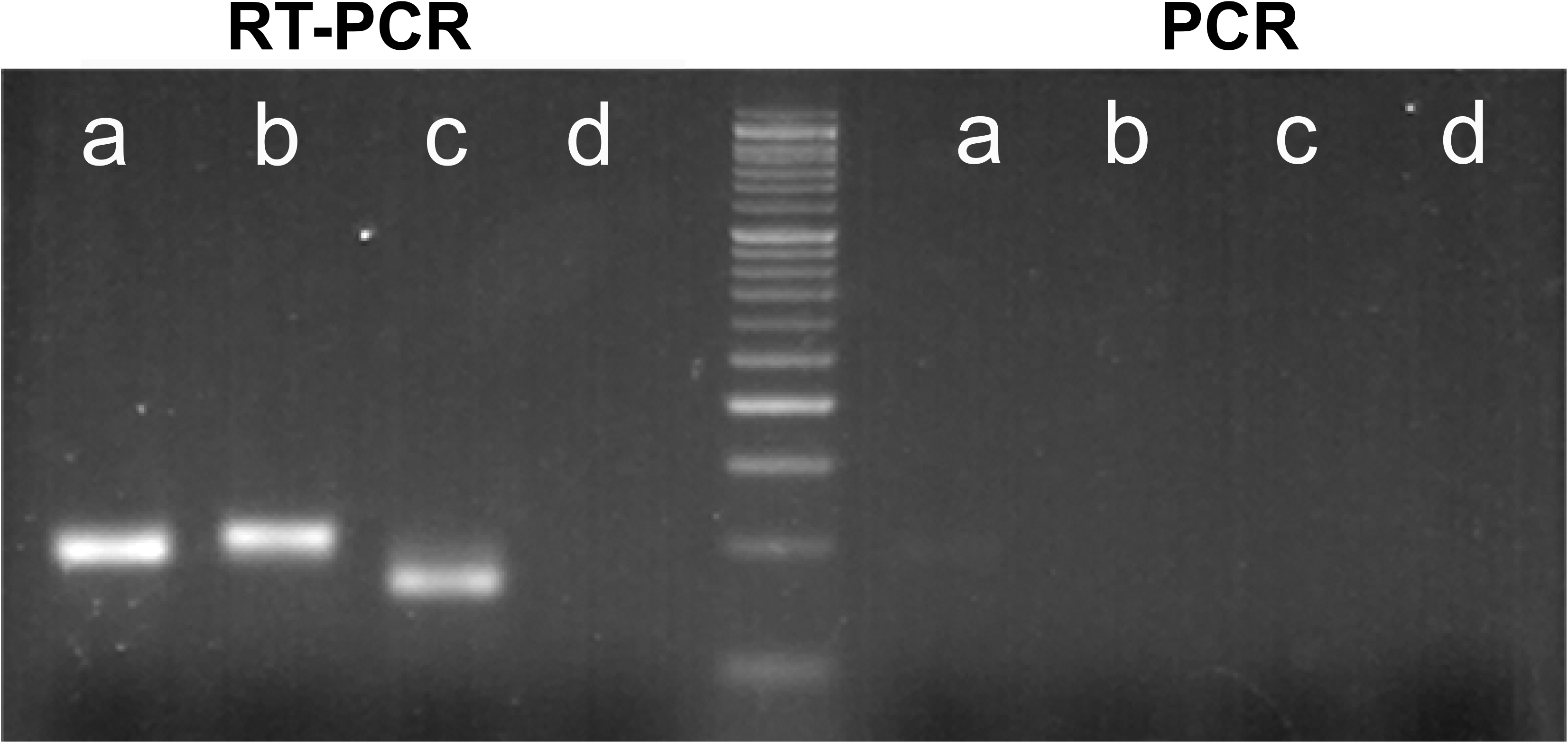
RT-PCR and PCR for confirmation of IVT RNA integrity. Specific amplification from IVT RNA was achieved by RT PCR while there was no amplification by PCR which confirms the lack of carry-over DNA template in IVT RNA. Lane a - MERS-CoV *UpE* (92 bp), lane b – NiV *N* (100 bp), lane c - REBOV *Vp40* (80 bp), lane d-NTC. Ladder - 50 bp.

### Optimization of hapten labels and antibody for NALFIA device

The hapten labels DIG, fluorescein, texas red, alexa fluor 488 and rhodamine red reacted with their corresponding immobilized antibodies on the nitrocellulose lateral flow strip. The hapten label DNP did not show any working signal with its corresponding immobilized antibody line on the lateral flow strip. There was non-specific capture of DIG label by immobilized polyclonal anti-fluorescein and polyclonal anti-texas red antibody lines. The non-specific capture of DIG by polyclonal anti-fluorescein antibody was not seen when the polyclonal anti-fluorescein antibody line was replaced by monoclonal anti-fluorescein antibody. There was cross-reactivity between rhodamine red and texas red labels and their antibodies. The reactivities of labels to their corresponding antibodies and the non-specific and cross-reactivities for multiplexing compatibility are summarized in (Table 4). Out of the six hapten-label combinations, the finalized 5’ labels on forward primers of NiV *N*, MERS-CoV *UpE* and REBOV *Vp40* for triplex NALFIA were digoxigenin, rhodamine red and alexa fluor 488. Fig 7 shows the signal development on hexaplex format and triplex format.

**Fig 7:**
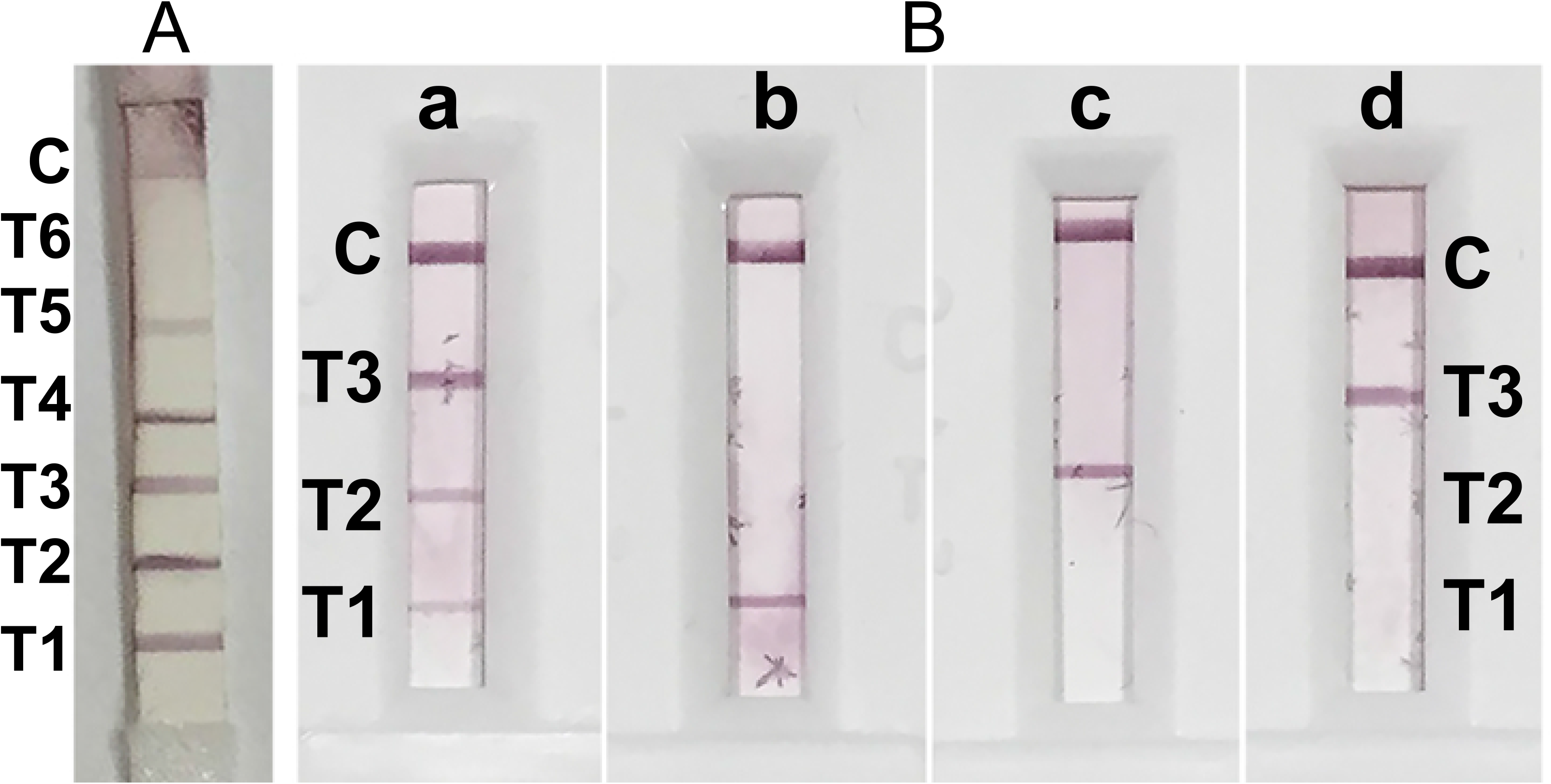
Optimisation of 5’ primer label/tag and antibody combinations for NALFIA. A - Hexaplex format; T6 - Dinitrophenyl (DNP), T5 - Digoxigenin (DIG), T4 - Fluorescein, T3 - Texas-red, T2 - Alexa fluor 488, T1 - Rhodamine. Out of the 6 label and antibody combinations tested, DNP and anti-DNP showed no reactivity while texas red antibody showed cross-reactivity with multiple labels. B - Triplex format; three labels DIG (T3), rhodamine red (T2) and alexa fluor 488 (T1) were finally selected and all further experiments were conducted using this combination.

**Table 4:**
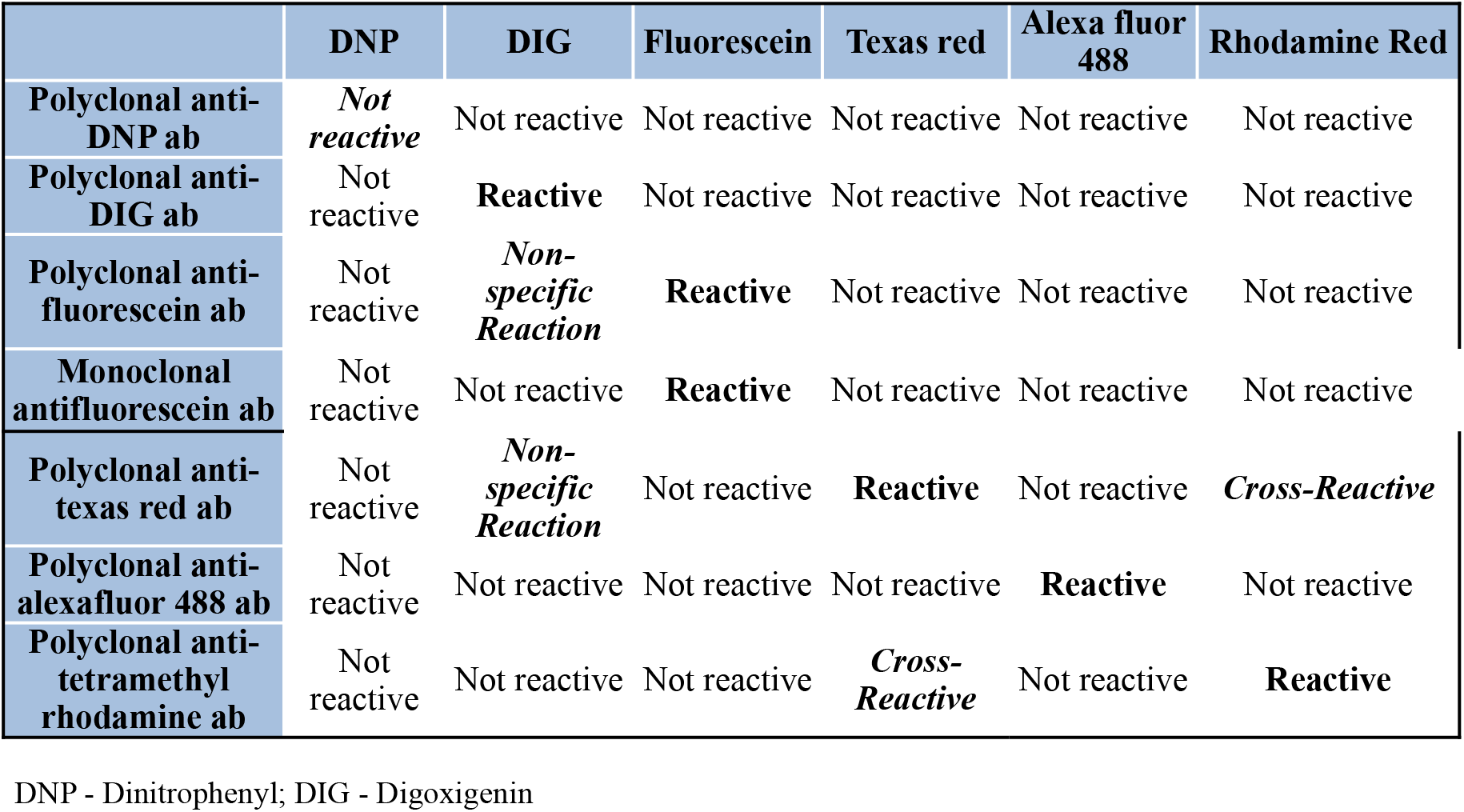
Summary of reactivity and compatibility evaluation of label and antibody combinations for multiplexing on NALFIA device

### Sensitivity of NALFIA device for labelled PCR amplicons

Two microliter of labelled PCR product was detectable on the NALFIA device till 10^-2^ dilution and the band intensity in NALFIA device reduced with the increase in dilution for all the three targets analyzed, which corresponded to the 3% agarose gel band gradation (Fig 8).

**Fig 8:**
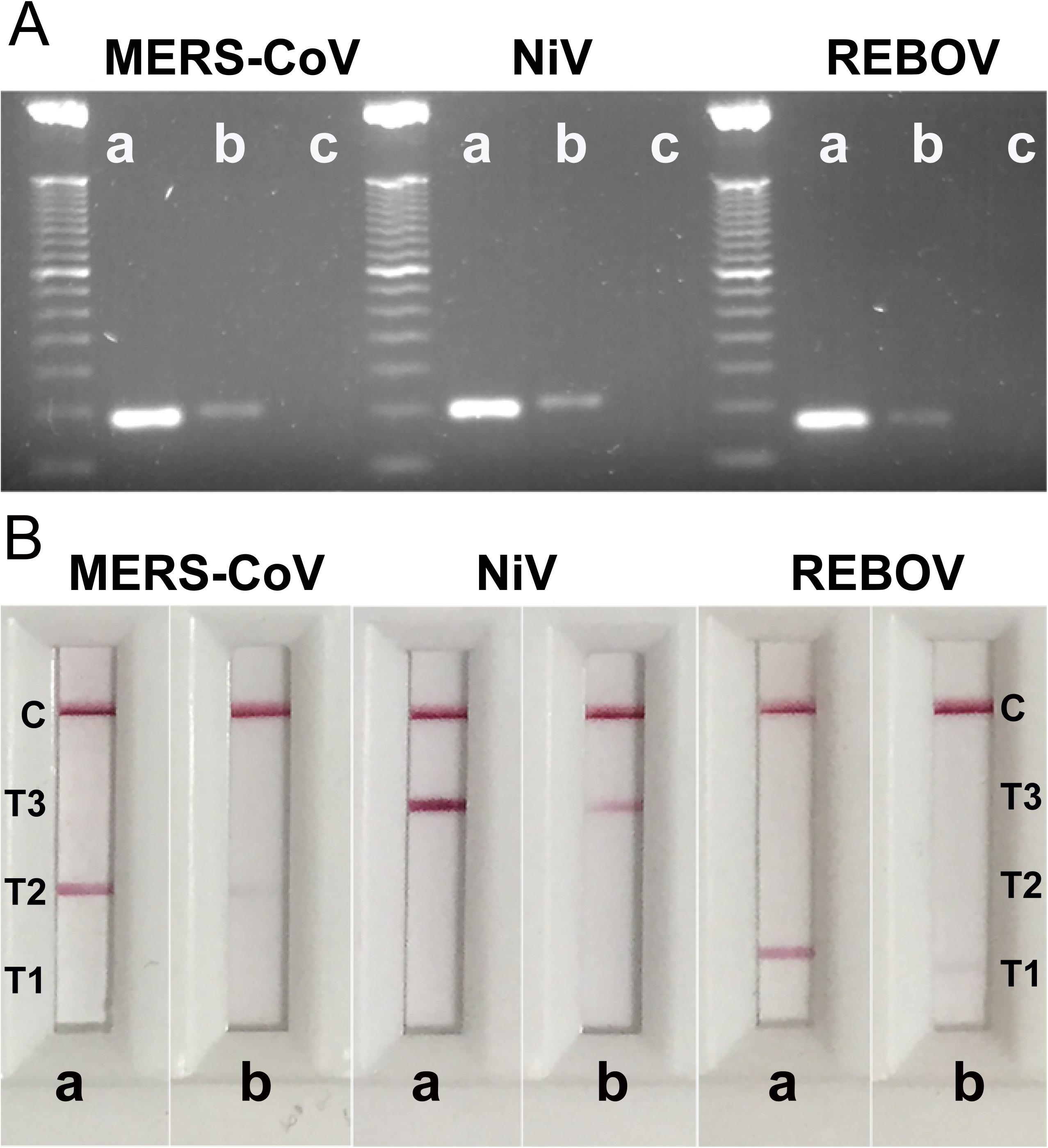
Sensitivity of NALFIA device for labelled PCR amplicons. A - Band intensity of 5 μl PCR product on 3% Agarose gel; lane a - undiluted PCR product, lane b - 1e-1 dilution, lane c - 1e-2, Ladder - 50 bp. B - Band intensity of 2 μl PCR product on NALFIA device; lane a - undiluted PCR product, lane b - 1e-1.

### Limit of detection (LOD) of Triplex NALFIA

The lowest copy numbers of IVT RNA that were detectable by triplex NALFIA with 2 μl of triplex RT-PCR products of 13 μl reaction were upto8.21 × 10^4^ for NiV *N* target, 7.09 × 10^1^ for MERS-CoV *UpE* target, and 1.83 x10^4^ for REBOV *Vp40* target (Fig 9). 3% agarose gel electrophoresis revealed that the RT-PCR amplification reduced linearly with the increase in IVT RNA dilution for all the three targets analyzed (S2).

**Fig 9:**
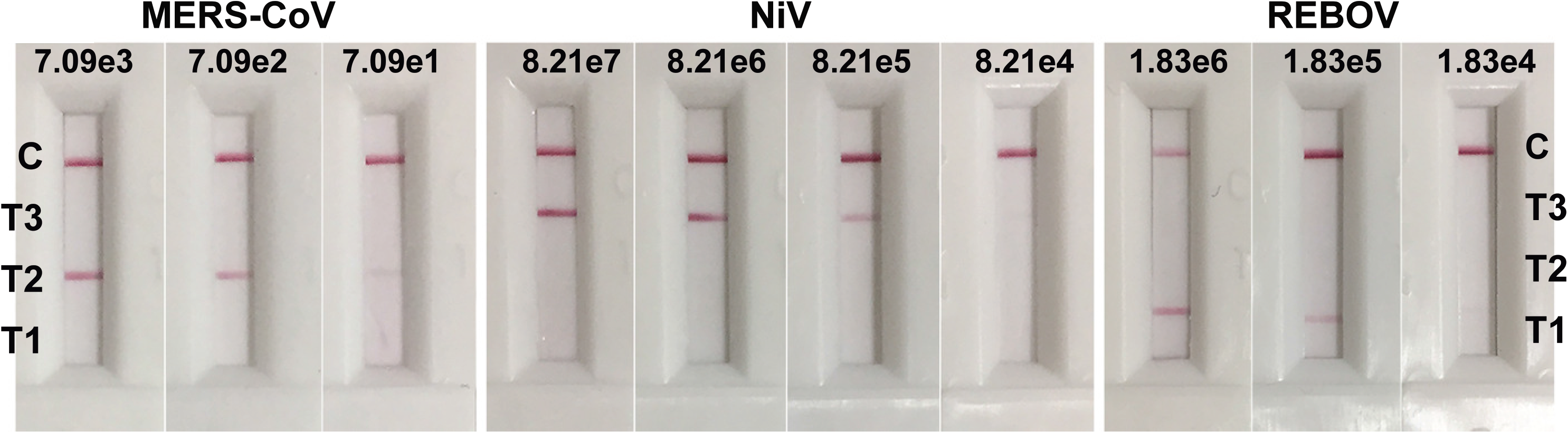
Analytical sensitivity of triplex-NALFIA. The IVT RNA was detectable up to 7.09e1 for MERS-CoV *UpE* target, 8.21e4 for NiV *N* target and 1.83e4 for REBOV *Vp40* target.

### Specificity of Triplex NALFIA

Band that corresponds to the specific target developed on the triplexed NALFIA device and no cross-amplification and cross-reactivity was observed between the three targets (Fig 9). Clinical samples positive for *Flaviviridae* (Japanese encephalitis virus, JEV; Classical swine fever virus, CSFV), *Paramyxoviridae* (New castle disease virus, NDV), *Coronaviridae* (Infectious bronchitis virus, IBV), *Orthomyxoviridae* (Avian influenza virus, AIV), *Picornaviridae* (Foot and mouth disease virus, FMDV),) *Arteriviridae* (Porcine reproductive and respiratory syndrome virus, PRRSV), *Herpesviridae* (Duck plague virus, DPV), *Poxviridae* (Swinepox virus, SPV) did not develop any visible signal on the bands designated for NiV, MERS-CoV and REBOV (Fig 10).

**Fig 10:**
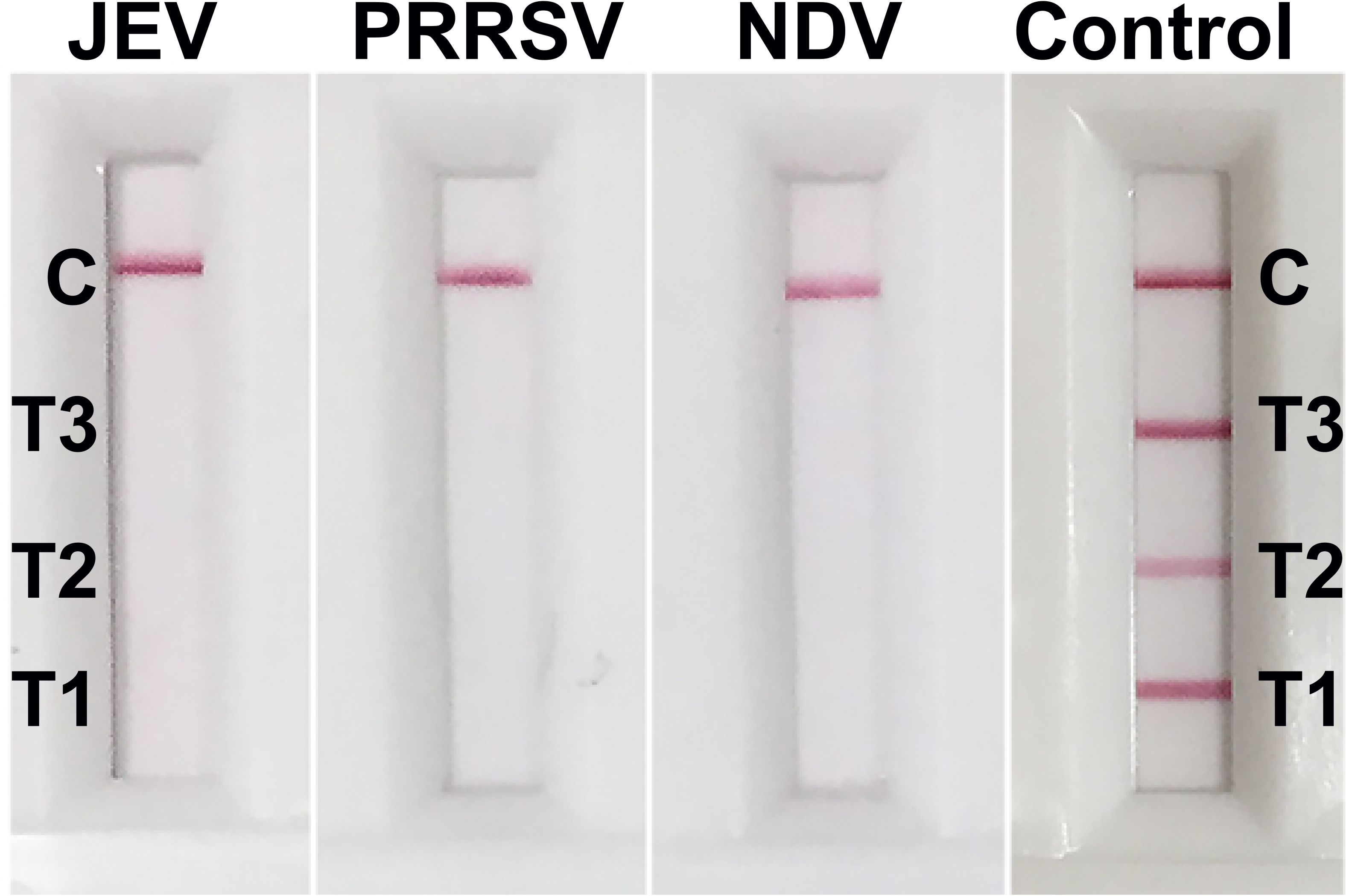
Triplex NALFIA negative for JEV, PRRSV and NDV.

### Evaluation of Triplex NALFIA with simulated samples

Eleven NiV *N* IVT RNA spiked samples and 15 un-spiked samples, 12 MERS-CoV *UpE* IVT RNA spiked samples and 18 un-spiked samples and 10 REBOV *Vp40* IVT RNA spiked samples and 14 un-spiked samples, verified by TaqMan RT-qPCR were tested by triplex NALFIA. Ten out of 11 NiV IVT RNA-positive samples were found to be positive by triplex NALFIA while 14 out of 15 negative samples were found to be negative by triplex NALFIA. All 12 MERS-CoV IVT RNA-positive samples were found to be positive by triplex NALFIA and all 18 negative samples were found to be negative by triplex NALFIA. All 10 REBOV IVT RNA-positive samples were found to be positive by triplex NALFIA and all 14 negative samples were found to be negative by triplex NALFIA. Also, no cross reactivity was observed between the targets. The sensitivity and positive predictive value were 91% for NiV *N* target and 100% for both MERS-CoV *UpE* and REBOV *Vp40* targets. The specificity and negative predictive value were 93.3% for NiV *N* target and 100% for both MERS-CoV *UpE* and REBOV *Vp40* targets. Overall sensitivity and specificity of the triplex NALFIA were 96% and 97%, respectively.

### Repeatability

The triplex NALFIA conducted three times on two simulated positive samples of each of the three targets all gave consistent positive results, hence indicating100% repeatability of the assay.

## Discussion

We have developed the first triplex NALFIA for the simultaneous detection of genomes of NiV, MERS-CoV and REBOV for screening bats and other respective hosts and reservoirs of these three viruses. Emerging infectious diseases (EIDs) are a significant threat to global public health, and bats are increasingly implicated in viral EID events by serving as reservoir hosts for viruses that can opportunistically cross species barriers to infect humans and other domestic as well as wild mammals. In the human outbreaks of Nipah virus (NiV), Middle East respiratory syndrome coronavirus (MERS-CoV), and ebolavirus, bats have been directly or indirectly implicated with or without the involvement of an intermediate host. For identifying the prevalence of these viruses in bats and other respective hosts and reservoirs, continuous monitoring and swift implementation of prevention and control policies, rapid and user-friendly diagnostics of these viruses are required. Assays available for virus detection and diagnosis mostly require centralized facility and sophisticated operation, which renders them unsuitable for routine screening in remote and low resource settings. Multiplexing saves time, allows testing with limited sample quantity, thus is suitable in low resource settings for simultaneous three-virus surveillance and first-line screening of NiV, MERS-CoV and REBOV infections in bats and other hosts. The main concern of a multiplex assay is the potential cross-reactivity, which limits the number and types of biomarkers that can be combined. Careful selection of primers and optimization of hapten-label combination are reported for successful multiplexing in amplification based NALFIA for other pathogens in previous studies (16, 18, 20, 29). Our work was conceptualized based on the need and benefits for a multiplexed assay in our region where multiple NiV outbreaks in human have been reported with evidences of prevalence in bats (30, 31, 32, 33) alongside threats of introduction of MERS-CoV and REBOV from neighboring countries. The triplex NALFIA, although intended for bats, can also be used for screening of other hosts of these viruses as it detects the virus genomes irrespective of the host.

The current assay reports the use of triplex one-step RT-PCR amplification with labelled forward and reverse primers and the visual detection of specific PCR amplicons on triplexed NALFIA device. Multiplexed NALFIA device with PCR-based amplification using labelled forward and reverse primers has been reported with high analytical sensitivity and specificity in a previous study for detection of Plasmodium DNA (19). In another study of multiplexed NALFIA for DNA detection and differentiation of three diarrhoeal protozoa, a labelled probe and a labelled primer, and an unlabelled primer were employed for isothermal amplification (16).

The sensitivity of the present NALFIA device for end point detection of the triplex RT-PCR product was compared with agarose gel electrophoresis and was found to correspond and even superior to the sensitivity of agarose gel electrophoresis which indicates that NALFIA is a rapid alternative to agarose gels. The analytical sensitivity of NALFIA device was even reported to be higher by 10-fold than the analytical sensitivity of agarose gel electrophoresis in a previous report (29). NALFIA device as naked eye end-point detection method not only overcomes the complexity of conventional post-amplification steps but also introduces a scope of point-of-care in the nucleic acid amplification-based diagnosis of infectious diseases. It is evident from previous reports on detection of other organisms that the isothermal amplification backed NALFIA presents itself to be a good option for POCD (16, 34, 35). Recently, LAMP and other isothermal assay-based multiplex NALFIA has been successfully developed for detection of multiple gene targets of SARS-CoV 2 demonstrating the feasibility of multiplexed NALFIA (21, 22, 23).

The primer sequences used in the present study are reported primers that have been validated in previous real-time RT-PCR assay development studies. This circumvents the essential validation on different organisms. Adaptation of published primers in other assays has been reported (36). Cross-amplification was analyzed at the level of RT-PCR and cross-reactivity at end-point on NALFIA device. Amplification was specific for the respective selected targets and no cross amplification was observed. At the level of NALFIA device, six combinations of label and antibody were initially tested for specific reactivity and compatibility in multiplexing. Haptens for 5’ end labeling of forward oligos were dinitrophenyl (DNP), digoxigenin (DIG), texas red, fluorescein, rhodamine red and alexa fluor 488. The antibodies were polyclonal anti-DNP, polyclonal anti-DIG, polyclonal anti-texas red, polyclonal anti-fluorescein, monoclonal anti-fluorescein, polyclonal anti-tetramethyl rhodamine, and polyclonal anti-alexa fluor 488. Out of the six tested; there was no working signal at the DNP test line. The failed interaction between DNP and its antibody could be due to various factors–stearic hindrance, low sensitivity of antibody, batch and lot defect etc. Further testing of the combination by changing the brand and the lot of both DNP and its antibody could give a successful combination as successful use of DNP was already reported in a previous assay (18). However, due to time constraints, the DNP combination was not tested further and was excluded from the subsequent assays. There was cross-reactivity between texas red and rhodamine red which could be due to structural relatedness between the two compounds. Although not related, the polyclonal anti-texas red antibody also reacted non-specifically with DIG label. The DIG label was also found to react non-specifically with the polyclonal anti-fluorescein antibody but this problem was solved when polyclonal antibody was replaced by monoclonal antibody. Similarly, the problem of non-specific reactivity seen with anti-texas red antibody with DIG label could also be solved by using monoclonal antibody; however, this experiment was not performed due to high cost involved with monoclonal antibody. Subsequently, the three final label-antibody combinations selected for our triplex-NALFIA were DIG, rhodamine red, alexa fluor 488 and their corresponding antibodies mentioned above. Alexa fluor 488 and rhodamine red have already been successfully employed in a previous study of a prototype multiplex NALFIA (16). DIG has been used as a standard label in most monoplex and duplex NALF assays (18, 20, 37). Testing of other label antibody combinations have also been discussed briefly in a previous study (16). The three final label-antibody combinations selected in the present study were not cross-reactive. However, out of the three label-antibody combinations, the sensitivity of the combination of rhodamine red and anti-tetramethyl rhodamine antibody was found to be slightly lower than that of the other two, as revealed by the weaker signal at MERS-CoV test line on the NALFIA device despite the amplification product having similar band intensity on AGE as the other two targets. The reading was recorded within a maximum time of 15 minutes and appearance of non-specific and ghost bands observed beyond this period were ignored. Most reported NALFIA are monoplex or duplex and the reports on multiplex NALFIA are limited. Therefore, the present work is an attempt to demonstrate the compatibility of the tags in a multiplex set-up which are otherwise successfully employed in a monoplex setup. Our hapten label streamlining strategy is novel although slightly similar toa previous work by Crannel and his team (16), and the current assay was conceptualized independently as the said previous report was not published at the time of development of the current assay.

The triplex NALFIA was evaluated on negative samples positive for Japanese encephalitis virus, classical swine fever virus New Castle disease virus, infectious bronchitis virus, avian influenza virus, foot and mouth disease virus, porcine reproductive and respiratory syndrome virus, suipox virus, and duck plaque virus, however, no signal was developed which indicates the specificity of the assay for its intended targets. Also, dimers that could have been generated during amplification did not produce visible signal on NALFIA device. Due to unavailability of clinically positive samples at the time of development of the current assay, a simulation of positive samples was created by spiking samples comprised of sera, swab, tissues and blood from apparently healthy bat, pig and camel with IVT RNAs of the respective targets and the spiked samples were then used for evaluating the triplex NALFIA. Despite the same IVT RNA spiking concentration used, the positive response signal by triplex NALFIA varied from sample to sample which may be due to differences in final IVT RNA concentration post extraction. Nevertheless, there is a need for further evaluation of the operating range of the triplex NALFIA using quantitated virus, individually for each virus target within the corresponding sample matrix - spiked or natural infection. However, this can only be performed at a BSL4 containment facility. The label and antibody combination highlighted in the present assay can be applied to detection of other pathogens if the primers are replaced with specific primers corresponding to desired targets. This is the first report on NALFIA based molecular detection tool for NiV, MERS-CoV and REBOV.

## Acknowledgement

We thank the Department of Biotechnology, Ministry of Science and Technology, Government of India (DBT) for partially funding this work under grant number ADMaC DBT-NER/LIVS/11/2012. We thank the UBio Biotechnology Ltd, Kochi (India) for the fabrication of NALFIA device. We also thank the Director, ICAR-National Institute of High Security Animal Diseases (NIHSAD), Bhopal, India.

The authors declare no competing interests.

